# Changes in the senescence profile and immune checkpoints in HIV-infected individuals after COVID-19

**DOI:** 10.1101/2024.03.12.584682

**Authors:** Celia Crespo-Bermejo, Óscar Brochado-Kith, Sergio Grande-García, Violeta Lara-Aguilar, Manuel Llamas-Adán, Sonia Arca-Lafuente, Luz Martín-Carbonero, Ignacio de los Santos, M Ángeles Jiménez Sousa, Salvador Resino, Juan Berenguer, Ricardo Madrid, Amanda Fernández-Rodríguez, Verónica Briz

## Abstract

**Background:** Both SARS-CoV-2 and HIV infection exhibit alterations in the senescence profile and immune checkpoint (IC) molecules. However, the midterm impact of SARS-CoV-2 on these profiles in people with HIV (PWH) remains unclear. This study aimed to evaluate differences in plasma biomarker levels related to ICs, the senescence-associated secretory phenotype (SASP), and pro- and anti-inflammatory cytokines in PWH following recovery from SARS-CoV-2 infection.

**Methods:** We conducted a cross-sectional study of 95 PWH receiving antiretroviral therapy, stratified by SARS-CoV-2 infection status: a) 48 previously infected (HIV/SARS) and b) 47 controls without previous infection (HIV). Plasma biomarkers (n=44) were assessed using Procartaplex Multiplex Immunoassays. Differences were analyzed using a generalized linear model adjusted for sex and ethnicity and corrected for the false discovery rate. Significant values were defined as an adjusted arithmetic mean ratio ≥1.2 or ≤0.8 and a qvalue<0.1. Spearman correlation evaluated relationships between plasma biomarkers (significant correlations, rho≥0.3 and q value<0.1).

**Results:** The median age of the PWH was 45 years, and 80% were men. All SARS-CoV-2-infected PWH experienced symptomatic infection; 83.3% had mild symptomatic infection, and sample collection occurred at a median of 12 weeks postdiagnosis. The HIV/SARS group showed higher levels of ICs (CD80, PDCD1LG2, CD276, PDCD1, CD47, HAVCR2, TIMD4, TNFRSF9, TNFRSF18, and TNFRSF14), SASP (LTA, CXCL8, and IL13), and inflammatory plasma biomarkers (IL4, IL12B, IL17A, CCL3, CCL4, and INF1A) than did the HIV group.

**Conclusions:** SARS-CoV-2 infection in PWH causes significant midterm disruptions in plasma ICs and inflammatory cytokine levels, highlighting SASP-related factors, which could be risk factors for the emergence of complications in PWH.

## 1. INTRODUCTION

The coronavirus pandemic has modified the quality of life of the general population, and although severe acute respiratory syndrome coronavirus 2 (SARS-CoV-2) causes acute infection to develop, follow-up studies have revealed that mid and long-term COVID-19 in the general population is associated with systemic effects, neurological issues, cardiovascular abnormalities, and pulmonary sequelae [1–3]. The impact of SARS-CoV-2 infection on the immune and inflammatory systems is particularly concerning in people with HIV (PWH) due to their immunocompromised state and chronic inflammation, even when receiving ART [4]. However, the risk of acquiring SARS-CoV-2 in PWH compared to that in the general population is controversial [5, 6]. However, HIV has been identified as an independent predictor of long-term COVID-19 [7].

Previous studies have reported an exacerbation of the proinflammatory phenotype known as the senescence-associated secretory phenotype (SASP) due to SARS-CoV-2 infection [8], similar to what has been observed in PWH [9]. Moreover, SARS-CoV-2 infection may be a potential factor contributing to the onset and reappearance of different types of tumors [10–12].

HIV infection itself results in a heightened occurrence of either AIDS or non-AIDS-related cancer compared to that in general population [13, 14]. HIV infection leads to T-cell exhaustion because of persistent immune activation [15, 16]. The intricate balance between T-cell activation and autoimmunity involves a network of receptors known as immune checkpoint (IC) molecules [15], among others. The plasma levels of soluble immune checkpoint (sIC) molecules play a crucial role in and are linked to the onset and progression of different tumors. Indeed, they serve as potential biomarkers for the diagnosis and prognosis of cancer development and for assessing therapeutic responses [17–19].

In the context of SARS-CoV-2 [20, 21] and HIV infection [22, 23], both ICs and their soluble form [23] are upregulated, as is the SASP profile [8]. However, it remains unclear whether these ICs and SASP profiles are altered after COVID-19 resolution.

Currently, there is limited information assessing the midterm consequences of SARS-CoV-2 infection in PWH. Kolossváry et al. (2023) [24] revealed alterations in proteins linked to inflammatory and immune pathways after 3 months of follow-up. After a median follow-up of four and six months, Peluso and colleagues (2022) [25] and Mazzitelli et al. (2022) [26], respectively, demonstrated that PWH who had recovered from SARS-CoV-2 infection reported asthenia and neurocognitive symptoms. Additionally, Peluso et al. (2023) [27] suggested that PWH may be vulnerable to developing long COVID-19 or postacute COVID-19 syndrome (PACS). However, the impact of SARS-CoV-2 infection in PWH on the SASP and ICs has not been determined.

Thus, this study aimed to assess the midterm effects of SARS-CoV-2 infection on plasma biomarkers related to the SASP and immune-oncology checkpoints in controlled PWH under ART.

## 2. MATERIALS AND METHODS

### 2.1. Subjects

A cross-sectional study of plasma biomarkers was carried out in 95 PWH on ART stratified by SARS-CoV-2 infection status: a) 48 PWH who had SARS-CoV-2 infection previously (≥ 4 weeks postinfection and diagnosis with PCR+) (HIV/SARS) were obtained from the AIDS Research Network Cohort (CORIS) with plasma samples collected between April 1, 2020, and September 30, 2020. b) 47 PWH without previous SARS-CoV-2 infection (HIV) were used as a control group and were recruited before the onset of the COVID-19 pandemic and plasma specimens were stored at the National Center for Microbiology, Instituto de Salud Carlos III. The control and study groups were matched based on age and the onset of their initial symptoms, and none were vaccinated against SARS-CoV-2. The severity of COVID-19 was classified based on criteria established by the National Institutes of Health (NIH) [28].

In line with CDC guidelines, which state that long COVID-19 can manifest at least four weeks post-infection [29] and considering the follow-up data from previous studies [1, 30], we defined midterm effects as the impact of SARS-CoV-2 from at least 4 weeks up to a period of 6 months after the diagnosis of the infection.

Inclusion criteria for all participants in this study involved being older than 18 years and receiving successful antiretroviral treatment for undetectable viremia (viral load <50 copies/mL) for at least 1 year before sample collection. Exclusion criteria included pregnancy, the presence of antibodies or antigens against hepatitis B and C viruses, opportunistic infections, and other diseases such as neoplasms and cardiovascular and autoimmune disorders. The study adhered to the Declaration of Helsinki, and written consent was obtained from all PWH before enrollment. The Ethics Committee of Hospital General Universitario Gregorio Marañón approved the study (Internal Ref# 162/20), as did the Institute of Health Carlos III (CEI PI 18_2021).

### 2.2. Preparation and isolation of plasma samples

Plasma samples were obtained by centrifuging of peripheral blood in EDTA tubes. The plasma samples were clarified by centrifugation and stored at -80°C until use.

### 2.3. Plasma biomarkers

Procartaplex Multiplex Immunoassays (xMAP-Luminex Technology) (Thermo Fisher Scientific®) were used to quantify the plasma levels of 44 soluble biomarkers related to the SASP, inflammation, and cell IC molecules (**Table S1)**.

### 2.6. Statistical analysis

For the descriptive analysis of clinical and epidemiological data, we summarized categorical variables using frequency and percentage and continuous variables using the median and interquartile range (IQR). Significant differences between categorical data were calculated using the chi-squared test or Fisheŕs exact test when appropriate. The Mann-Whitney-Wilcoxon test was used to compare continuous variables among independent groups. Multivariate analysis with generalized linear models (GLMs) and gamma distributions (log-links) were carried out to estimate differences in the levels of biomarkers related to IC molecules, senescence analytes, and inflammatory cytokines among the groups. All tests were adjusted for sex and ethnic origin. P values were corrected by the false discovery rate (FDR) using the Benjamin-Hochberg correction, setting a cut-off point of 0.1. Significant values were defined as adjusted arithmetic mean ratio (aAMR) ≥1.2 or ≤0.8 and a q value<0.1. The relationship between plasma biomarkers was analyzed using Spearman correlation. Significant correlations were defined as a correlation coefficient (rho) ≥0.3 and a q value <0.1.

The statistical software R (v. 4.0.5) was used for all the statistical analyses.

## 3. RESULTS

We selected 47 PWH from individuals previously infected with SARS-CoV-2 and 48 controls.

### 3.1. Epidemiological and clinical characteristics of the patients

The sociodemographic and clinical characteristics of the study population are shown in **Table 1**. Overall, the patients had a median age of 45 years, 81.9% were Caucasian, and 80% were men. All PWH infected with SARS-CoV-2 reported symptoms. A total of 83.3% had mild COVID-19. The median interval between diagnosis and sample collection was 12 [IQR, 9-16] weeks.

**Table 1.** Sociodemographic and clinical characteristics of the study population. **Note: Statistics**: Values are expressed as the median [IQR] and absolute count (percentage). P-values were estimated by the Chi-square test for categorical variables and the Mann-Whitney-Wilcoxon test for continuous variables. Significant values are shown in bold type and were defined as p-value<0.05. **Abbreviations:** PWH, people with HIV; BMI, body mass index; II: integrase inhibitors; PI, protease inhibitors; NRTI, nucleotide reverse transcriptase inhibitors; NNRTI, nonnucleotide reverse transcriptase inhibitors; HIV, human immunodeficiency virus; IDU, intravenous drug users. NA, not applicable; IQR, interquartile range.

|  | **Overall**<br>(n=95) | **HIV**<br>(n=47) | **HIV/SARS**<br>(n=48) | **p value** |
| --- | --- | --- | --- | --- |
| **Gender** (male) | 76 (80.0%) | 32 (68.1%) | 44 (91.7%) | **0.005** |
| **Age** (years) | 45.2<br>[36.4-53.6] | 45.0<br>[36.0-54.0] | 45.4<br>[37.3-52.6] | 0.973 |
| **BMI** (kg/m<sup>2</sup>) | 24.9<br>[23.0-27.5] | 24.3<br>[22.7-26.0] | 25.9<br>[23.2-28.6] | 0.148 |
| **Ethnic origin**<br>(Caucasians) | 77 (81.9%) | 45 (97.8%) | 32 (66.7%) | **<0.001** |
| **HIV infection** |  |  |  |  |
| **Time of HIV infection**<br>(years) | 9.0<br>[5.0-13.0] | 9.0<br>[5.0-20.0] | 8.5<br>[5.0-12.0] | 0.311 |

| HIV antiretroviral therapy |  |  |  |  |
| --- | --- | --- | --- | --- |
| 2NRTI+II | 53 (56.4%) | 27 (57.4%) | 26 (55.3%) | 0.293 |
| 2NRTI+PI | 13 (13.7%) | 7 (14.9%) | 6 (12.5%) |  |
| 3NRTI | 24 (25.5%) | 11 (23.4%) | 13 (27.7%) |  |
| 2NRTI+NNRTI | 5 (5.3%) | 2 (4.3%) | 3 (6.4%) |  |
| HIV transmission routes |  |  |  |  |
| IDU | 2 (2.7%) | 0 (0.0%) | 2 (4.3%) | 0.770 |
| Sexual | 71 (94.7%) | 28 (96.6%) | 43 (93.5%) |  |
| Both | 2 (2.7%) | 1 (3.4%) | 1 (2.2%) |  |
| SARS-CoV-2 infection |  |  |  |  |
| COVID-19 severity |  |  |  |  |
| Mild | 40 (83.3%) | NA | 40 (83.3%) | NA |
| Severe | 8 (16.7%) | NA | 8 (16.7%) | NA |
| Diagnosis-sample collection (weeks) | 12.0 [9-16] | NA | 12.0 [9-16] | NA |

### 3.2. SASP profile and other anti and proinflammatory cytokines in PWH after recovering from SARS-CoV-2 infection

The HIV/SARS group presented significantly greater values for 3 out of the 13 cytokines related to the SASP profile, including LTA [aAMR= 1.36[1.03-1.80], q= 0.076], CXCL8 [aAMR= 1.41[1.13-1.76], q= 0.020] and IL13 [aAMR= 1.30[1.04-1.62], q= 0.076]. In addition, the HIV/SARS group exhibited significantly greater alterations in the expression of 6 out of the 8 anti- and proinflammatory cytokines, including IL4 [aAMR= 1.27[1.03-1.55], q= 0.076], IL12B [aAMR= 1.57[1.21-2.03], q= 0.011], IL17A [aAMR= 1.30[1.06-1.59], q= 0.038], CCL3 [aAMR= 2.60[1.47-4.61], q= 0.011], CCL4 [aAMR= 1.25[1.01-1.56], q= 0.098], and IFNA1 [aAMR= 1.19[1.02-1.40], q= 0.076]. (**Figure 1**) (**Tables S2 and S3)**.

**Figure 1.**
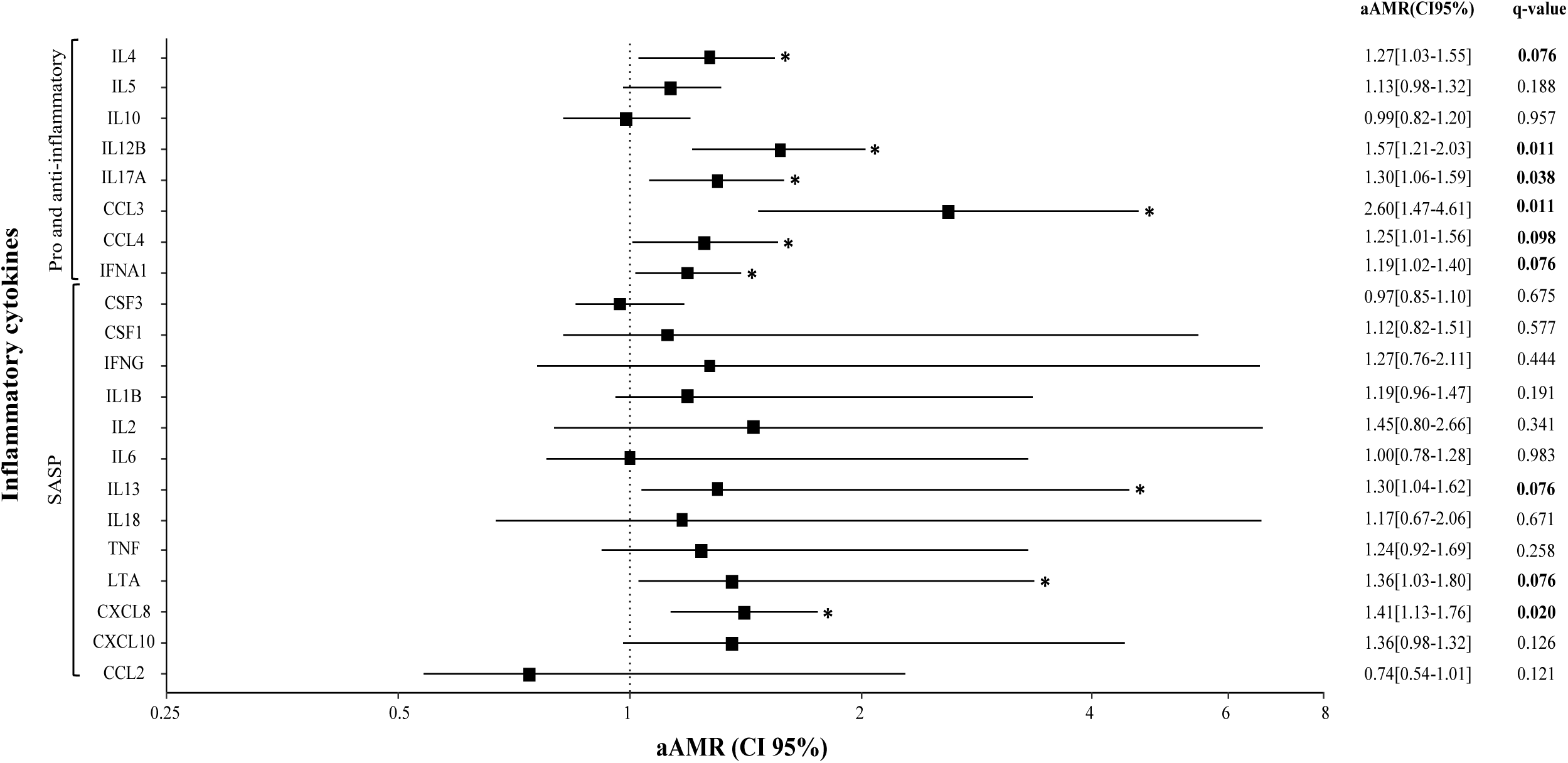
Association of the SASP and pro- and anti-inflammatory cytokines levels in plasma with midterm effects of SARS-CoV-2 infection in PWH. Forest plot representing the aAMR and 95% CI of the SASP and pro- and anti-inflammatory plasma cytokines in PWH after recovering of SARS-CoV-2 infection. Statistics: Differences between groups were analyzed using a GLM, adjusted for sex and etnia. *Significant values were defined as aAMR≥1.2 or ≤0.8 and a q value<0.1. Abbreviations: aAMR, adjusted arithmetic mean ratio; CI, confidence intervals; q value, p value adjusted by multiple comparisons with the Benjamini and Hochberg correction.

### 3.3. Plasma immune checkpoint-related biomarkers in PWH after recovering from SARS-CoV-2 infection

The HIV/SARS group exhibited significantly greater alterations in 10 out of the 23 checkpoint biomarkers assessed (10/23) including: I) immune checkpoint biomarkers related to the immunoglobulin superfamily: CD80 [aAMR = 1.80 [1.27-2.56], q=0.011], PDCD1LG2 [aAMR = 1.25 [1.06-1.48], q=0.038], CD276 [aAMR = 1.28 [1.13-1.45], q=0.011], PDCD1 [aAMR = 1.65 [1.24-2.20], q=0.011] and CD47 [aAMR = 1.25[1.08-1.46], q=0.023], II) immune checkpoint biomarkers associated with the Tim family HAVCR2 [aAMR = 1.25 [1.10-1.42], q=0.011] and TIMD4 [aAMR = 1.37 [1.03-1.82], q=0.078], III) immune checkpoint biomarkers of TNF/RSF family TNFRSF9 [aAMR = 1.39 [1.03-1.87], q=0.082], TNFRSF18 [aAMR= 1.66 [1.16-2.37], q=0.033] and TNFRSF14 [aAMR = 1.35 [1.09-1.69], q=0.033](**Figure 2**) (**Table S4)**.

**Figure 2.**
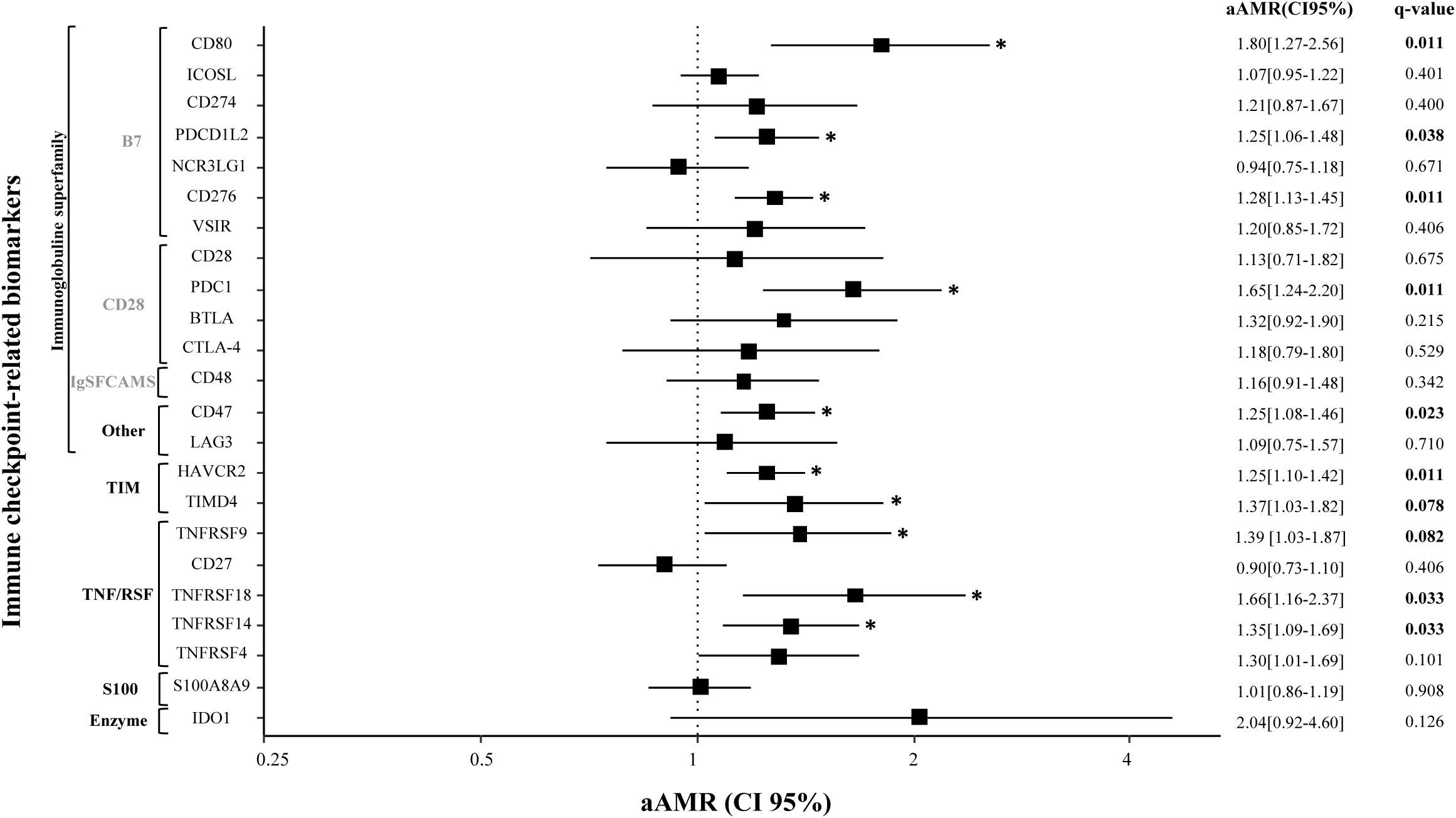
**Association of immune checkpoint-related biomarkers in plasma with midterm effects of SARS-CoV-2 infection in PWH**. Forest plot showing the aAMR and 95% CI for different checkpoint-related molecules. Statistics: Differences between groups were analysed using a GLM, adjusted for sex and ethnic origin, *Significant values were defined as aAMR≥1.2 or ≤0.8 and q value<0.1. Abbreviations: aAMR, adjusted arithmetic mean ratio; CI, confidence intervals; q value, p value corrected for multiple comparisons by Benjamini and Hochberg.

### 3.5. Correlations between significant plasma biomarkers in HIV/SARS patients

Significant positive correlations were found between inflammatory cytokines, more specifically between the levels of the SASP cytokines IL13 and LTA, and the levels of the IC molecules CD80 [rho=0.31, q=0.073; rho=0.34, q=0.073, respectively], TNFRSF18 [rho=0.37, q=0.024; rho=0.44, q=0.019, respectively], and TNFRSF14 [rho=0.33, q=0.071; rho=0.46, q=0.012, respectively] (**Figure 3**) (**Table S5)**.

**Figure 3.**
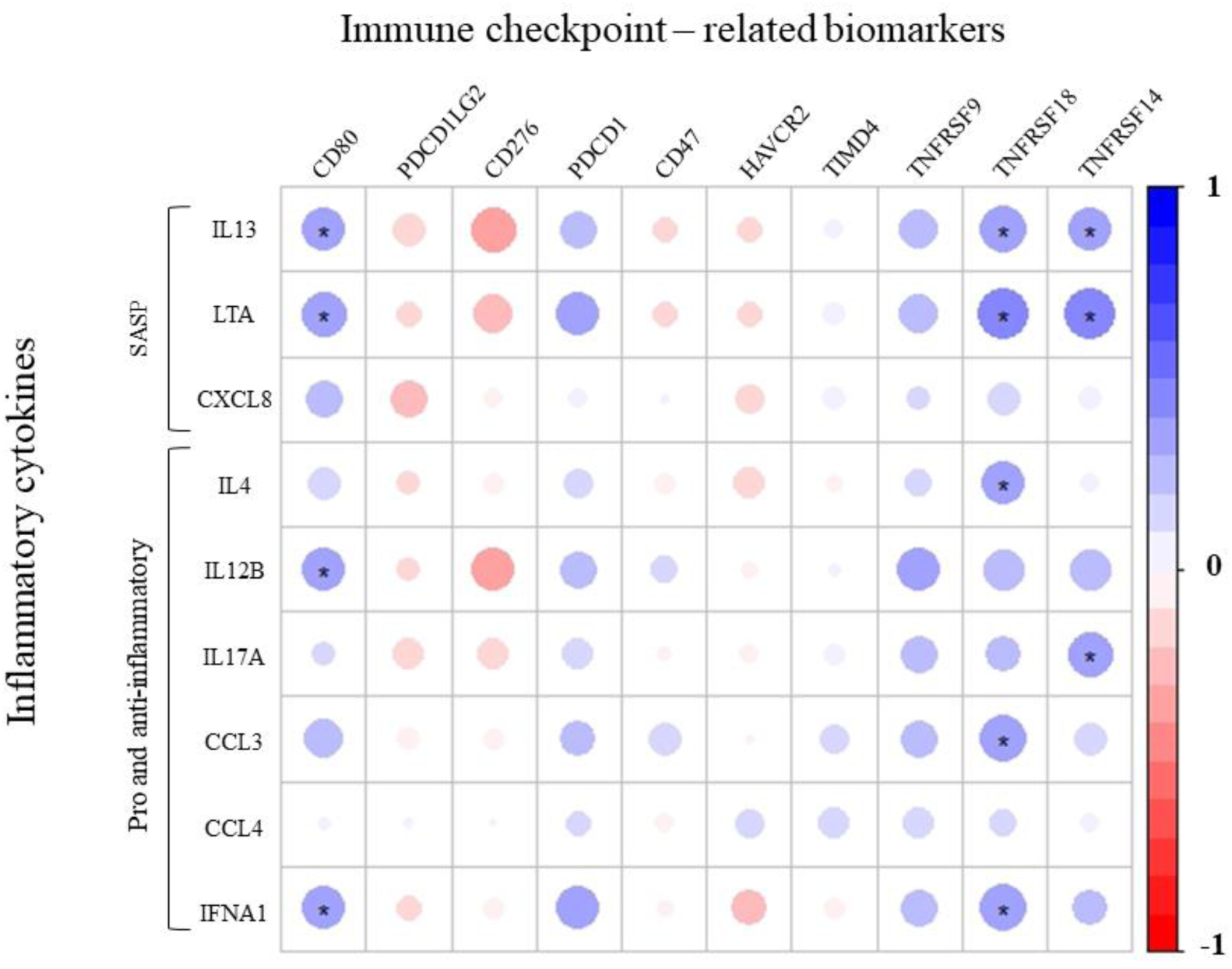
Relationships between significant plasma biomarkers in the HIV/SARS group. Correlation matrix showing the correlation between immune checkpoint-related molecules and pro- and anti-inflammatory cytokines in HIV/SARS patients. The size of the circles is proportional to the strength of the correlation, and the color represents the direction, where a large dark blue represents a strong positive correlation, and a large dark red circle represents a strong negative correlation. **Statistics**: analysis was performed with Spearman correlation. *Significant values were defined as rho≥0.3 and q value<0.1.

## 4. DISCUSSION

The midterm impact of SARS-CoV-2 infection in PWH showed significantly greater levels of the SASP-profile cytokines CXCL8, IL13, and LTA, as well as other anti- and proinflammatory cytokines, including IL4, IL12B, IL17A, CCL3, CCL4, and INFG, than in PWHs without previous SARS-CoV-2 infection. Additionally, midterm effects of SARS-CoV-2 infection were observed in biomarkers related to IC molecules, as indicated by significantly increased levels of sIC receptors and ligands, including CD80, PDCD1LG2, CD276 (B7 family), PDCD1, (CD28 family), CD47 (immunoglobulin superfamily) HAVCR2 and TIMD4 (TIM family), TNFRSF9, TNFRSF18, and TNFRSF14 (TNF/RSF family).

Our results are in accordance with those reported by Kolossváry et al. (2023), who also observed a significant dysregulation of proteins associated with inflammatory and immune pathways, such as CCL18 and CCL23, in PWH after 4 months of SARS-CoV-2 infection [24], compared to PWH without previous SARS-CoV-2 infection. However, our study identified distinct proteins, including 6 inflammatory cytokines (IL4, IL12B, IL17A, CCL3, CCL4, and IFNA1) and 3 SASP cytokines (CXCL8, IL13, and LTA) after midterm (12 weeks) SARS-CoV-2 infection. Differences in the selected cohort, as well as in the methodology used, could explain the variations in the results. On the one hand, all PWH included in our study had an undetectable viral load (VL) (<50 copies/ ml) for at least one year before sample collection. However, in Kolossváry’s cohort [24], 10% of PWH displayed virological failure [31] (400 copies/ml), which may result in ongoing immune activation and inflammation [32] and an additional source of cytokine disruption. On the other hand, all PWH included in Kolossváry’s study were collected between May 2020 and September 2021 based on any clinical diagnosis of COVID-19 and/or a positive SARS-CoV-2 rapid antigen detection test or PCR. The positivity of the samples was subsequently confirmed through an antibody test, but the absence of a diagnostic PCR+ test may introduce significant bias in estimating the follow-up date. Finally, the different methodologies used in both studies, the Luminex multiplex assay (ProcartaPlex Immunoassay) in our study and the proximity extension assay (PEA) technology (Olink®) [33], which identifies proteins using pairs of antibodies conjugated to cDNA strands, may also have influenced the final results. Additionally, 16 weeks after SARS-CoV-2 infection, the levels of the inflammatory cytokines IL6, CXCL10, and TNF were greater in PWH than in the general population [25]. The significant biomarkers were different from those identified in our study. Nevertheless, unlike our group, which comprises entirely PWH, they assessed seropositive vs. seronegative patients in this study. Increased levels of CXCL10 and TNF have been related to a higher risk of developing postacute sequelae of SARS-CoV-2 infection (PASC) in PWH [25].

To the best of our knowledge, only a limited number of studies have evaluated these plasma biomarkers in PWH after SARS-CoV-2 infection resolution [24, 25]. However, similar studies have been performed in the general population, where normalization of the serum levels of the inflammatory cytokines IL2, IL4, IL10, TNF, and IFNG [34] was observed after three weeks of SARS-CoV-2 resolution. However, contradictory results were shown by Ren and colleagues (2023) and Loretelli et al. (2021), as increasing circulating levels of the SASP-related cytokine CXCL8 were observed after 4.5 months [35] and 1 year [36] of SARS-CoV-2 infection. In addition, a recent report described IL12B and IL13 as altered cytokines 3 months after resolution of SARS-CoV-2 infection [37].

Our study also revealed increased levels of sIC receptors and ligands, such as members of the immunoglobulin superfamily, TNFRSF, and TIM families. Specifically, 12 weeks after the diagnosis of SARS-CoV-2, the levels of B7 family members (CD80, PDCD1LG2 and CD276), the CD28 family (PDCD1) and other nonspecified family (CD47), the TIM family (HAVCR2 and TIMD4), and members of the TNFRSF family (TNFRSF9, TNFRSF18, and TNFRSF14), increased in PWH. The TNFRSF, CD28-B7, and TIM families of ICs and ligands, such as sPDCD1 or sCD274, are closely related to the T-cell response [25]. In line with our results, Peluso et al. (2022) observed increased expression of PDCD1 on CD4+ T cells in PWH 16 weeks after recovering from COVID-19, which also led to T-cell exhaustion [25]. Therefore, considering the compromised immune system of PWH, acute infection would impede the return of patients to normal sIC levels. Conversely, the study of Kolossváry et al. (2023) [24] did not reveal differences in the analyzed IC receptors and ligands 4 months after SARS-CoV-2 infection. As metioned above, differences in the methodology and the inclusion and exclusion criteria for patients in both studies might influence the different impacts of SARS-CoV-2 infection. For instance, Kolossváry et al. [24] included PWH with a VL >50 copies/ml, which could directly impact sIC levels, as increasing plasma sICs, including PD-1 and TIM-3, have been observed in viremic and untreated HIV infection [38,39].

An increase of sIC levels has been observed during acute SARS-CoV-2 infection in the general population [23, 40–42]. Several studies have also reported higher sIC levels after the infection period, as indicated by increases in CD274 and TIGIT biomarkers [43], which were observed 9 months after acute infection, as well as elevated plasma levels of sPDCD1 4.5 months after SARS-CoV-2 infection, compared to those in HCs [35].

Future studies with longer follow-up periods are needed to assess the normalization of these altered cytokine levels or their association with additional comorbidities. In this respect, compared with those of the general population, PWH exhibit an elevated inflammatory state resembling that of elderly individuals with accelerated aging [44–46], therefore, the long-term impact of SARS-CoV-2 infection could be markedly different in PWH. In addition, they also bear an increased burden of preexisting senescent cells, which may be intensified by SARS-CoV-2 virus-induced senescence [47,48]. This might contribute to the development of comorbidities and mortality, which are often associated with advanced age [49].

Finally, we explored the correlations between the plasma biomarkers with significant differences among the groups. We observed a positive correlation between molecules related to the T-cell response, specifically CD80, TNFRSF18, and TNFRSF14, and pro and anti-inflammatory cytokines, such as IL4, IL12B, and IFNA1, which is in line with the findings of Roe and colleagues (2021) [50]. These results may suggest the occurrence of remaining T-cell exhaustion in PWH after 12 weeks of SARS-CoV-2 infection diagnosis. An increase in sPDC1, sHAVCR2, sTNFRSF14, and sCD80 has been previously described as a prognostic marker for cancer development [51–53]. Although there are no studies evaluating the effects of these elevated levels following SARS infection, their upregulation may lead to a putative early onset of tumorigenesis.

Finally, several aspects should be taken into account to interpret our data correctly. This was a preliminary study with a limited sample size, which could have limited the possibility of finding additional statistically significant findings. However, despite our sample size, we identified meaningful distinctions within the dataset by applying adequate statistical methods and meticulous data collection. Another limitation that should be mentioned is the lack of plasma samples from the same patients before and after SARS-CoV-2 infection, which allows for a more comprehensive follow-up. Additionally, the inability to confirm negativity through PCR testing after recovery is another notable limitation. In fact, we applied FDR correction to limit false-positive results. Additional studies with longer follow-ups are necessary to analyze both the dynamics of IC molecules and the SASP and the occurrence of possible cancer-related events in PWH after COVID-19.

## 5. CONCLUSIONS

The impact of SARS-CoV-2 infection on PWH leads to the disruption of plasma biomarkers, such as sIC, the SASP, and pro- and anti-inflammatory cytokines, which increase at a median of 12 weeks after diagnosis. These results could lead to the development of risk factors for accelerating T-cell immune exhaustion and triggering potential future cancer-related events.

## 6. DECLARATIONS

### 6.1. Ethical Approval and Consent to participate

The study was approved by the Ethics Committee of Hospital General Universitario Gregorio Marañón (Internal Ref# 162/20) and the Institute of Health Carlos III (CEI PI 18_2021) and the study was performed in accordance with the ethical standards of the 1964 Declaration of Helsinki and its later amendments or comparable ethical standards. Informed consent was obtained from all patients included in the study.

### 6.2. Consent for publication

Not applicable.

### 6.3. Availability of data

The datasets used and/or analyzed in the present study are available from the corresponding author upon reasonable request.

### 6.4. Authors’ contributions and materials

Funding body: Verónica Briz and Ricardo Madrid. Study concept and design: Verónica Briz, Ricardo Madrid and Amanda Fernández-Rodríguez. Patient selection and clinical data acquisition: Luz Martín-Carbonero, Ignacio de los Santos, and Juan Berenguer. Sample preparation and analysis: Celia Crespo-Bermejo, Violeta Lara-Aguilar, Manuel Llamas-Adán, Sonia Arca-Lafuente, Sergio Grande-García. Statistical analysis and interpretation of the data: Celia Crespo-Bermejo, Óscar Brochado-Kith, Amanda Fernández-Rodríguez, and María Ángeles Jiménez-Sousa. Writing of the manuscript: Celia Crespo-Bermejo. Critical revision of the manuscript for relevant intellectual content: Verónica Briz, Amanda Fernández-Rodríguez and Ricardo Madrid. Supervision and visualization: Amanda Fernández-Rodríguez, Verónica Briz, Ricardo Madrid, Salvador Resino, María Ángeles Jiménez-Sousa, Luz Martín-Carbonero, and Ignacio de los Santos. All the authors read and approved the final manuscript.

## 6.6. Acknowledgments

The authors wish to thank all the patients and the nursing team for their participation in this study.

