## Supplemental data for "Changes in the senescence profile and immune checkpoints in HIV-infected individuals after COVID-19"

**Table S1. Procartaplex Multiplex Immunoassays (Thermo Fisher Scientific®) panels.**

- *Immuno-Oncology Checkpoint 14-Plex Human ProcartaPlex™ Panel 1*

| Marker Name | Official Symbol | Full description |
|---|---|---|
| **Immunoglobuline superfamily** | | |
| | ***B7 family*** | |
| CD80 | CD80 | CD80 molecule |
| PD-L1 | CD274 | CD274 molecule |
| PD-L2 | PDCD1LG2 | Programmed cell death 1 ligand 2 |
| | ***CD28 family*** | |
| CD28 | CD28 | CD28 molecule |
| PD1 | PDC1 | Programmed cell death 1 |
| BTLA | BTLA | B and T lymphocyte associated |
| CD152 (CTLA4) | CTLA4 | Cytotoxic T-lymphocyte-associated protein 4 |
| | ***Non-specific family*** | |
| LAG3 | LAG3 | Lymphocyte activating 3 |
| **TIM family** | | |
| TIM3 | HAVCR2 | Hepatitis A virus cellular receptor 2 |
| **TNF/RSF family** | | |
| CD137 (4-1BB) | TNFRSF9 | TNF receptor superfamily member 9 |
| CD27 | CD27 | CD27 molecule |
| GITR | TNFRSF18 | TNF receptor superfamily member 18 |
| HVEM | TNFRSF14 | TNF receptor superfamily member 14 |
| **Enzyme** | | |
| IDO1 | IDO1 | Indoleamine 2,3-dioxygenase 1 |

- *Immuno-Oncology Checkpoint 9-Plex Human ProcartaPlex™ Panel 3*

| Marker Name | Official Symbol | Full description |
|---|---|---|
| **Immunoglobuline superfamily** | | |
| | ***B7 family*** | |
| ICOSL (B7-H7) | ICOSL | Icos ligand |
| B7H6 | NCR3LG1 | Natural killer cell cytotoxicity receptor 3 ligand 1 |
| CD276 (B7H3) | CD276 | CD276 molecule |
| VISTA (B7-H5) | VSIR | V-set immunoregulatory receptor |
| | ***IgFCAMS family*** | |
| CD48 (BLAST-1) | CD48 | CD48 molecule |
| | ***Non-specific family*** | |
| CD47 (IAP) | CD47 | CD47 molecule |
| **TIM family** | | |
| TIMD4 | TIMD4 | T cell immunoglobulin and mucin domain containing 4 |
| **TNF/RSF family** | | |
| CD134 (OX40) | TNFRSF4 | TNF receptor superfamily member 4 |
| **S100** | | |
| S100-A8/A9 | S100-A8/A9 | S100 calcium binding protein A8/A9 |

- *Human Cytokine Storm Panel 21-plex*

| Target Name | Official Symbol | Full description |
|---|---|---|
| ***SASP-related cytokines*** | | |
| G-CSF (CSF-3) | CSF3 | Colony stimulating factor 3 |
| GM-CSF (CSF-1) | CSF2 | Colony stimulating factor 2 |
| IFN gamma | IFNG | Interferon gamma |
| IL-1β | IL1B | Interleukin 1 beta |
| IL-2 | IL2 | Interleukin 2 |
| IL-6 | IL6 | Interleukin 6 |
| IL-13 | IL13 | Interleukin 13 |
| IL-18 | IL18 | Interleukin 18 |
| TNF-α | TNF | Tumor necrosis factor alpha |
| TNF-β | LTA | Lymphotoxin alpha |
| IL-8 (CXCL8) | CXCL8 | Interleukin-8 |
| IP-10 (CXCL10) | CXCL10 | C-X-C motif chemokine ligand 10 |
| MCP-1 (CCL2) | CCL2 | C-C motif chemokine ligand 2 |
| ***Pro and anti-inflammatory cytokines*** | | |
| IL-4 | IL4 | Interleukin 4 |
| IL-5 | IL5 | Interleukin 5 |
| IL-10 | IL10 | Interleukin 10 |
| IL-12p70 | IL12B | Interleukin 12 |
| IL-17A (CTLA-8) | IL17A | Interleukin 17A |
| MIP-1 alpha (CCL3) | CCL3 | C-C motif chemokine ligand 3 |
| MIP-1 beta (CCL4) | CCL4 | C-C motif chemokine ligand 4 |
| IFN alpha | IFNA1 | Interferon Alpha 1 |

**Table S2. Levels of SASP-related cytokines on plasma in PLWH after recovering from SARS-CoV-2 infection.**

| | Mean (IQR) | | Univariate analysis | | | Multivariate analysis | | |
|---|---|---|---|---|---|---|---|---|
| **ANALITE** | **HIV** | **HIV/SARS** | **AMR** | **p** | **q** | aAMR | **p** | **q** |
| **Colony stimulating factors** | | | | | | | | |
| CSF3 | 17.00[15.50-19.00] | 16.00[14.00-19.00] | 0.99[0.89-1.10] | 0.843 | 0.862 | 0.97[0.85-1.10] | 0,614 | 0.675 |
| CSF1 | 8.00[7.00-10.50] | 10.25[8.00-13.00] | 1.16[0.89-5.51] | 0.259 | 0.317 | 1.12[0.82-1.51] | 0,472 | 0.577 |
| **Interferons** | | | | | | | | |
| IFNG | 22.00[17.00-39.00] | 29.00[19.88-71.75] | 1.34[0.88-2.06] | 0.174 | 0.233 | 1.27[0.76-2.11] | 0.343 | 0.444 |
| **Cytokines** | | | | | | | | |
| IL1B | 10.00[9.00-12.00] | 12.00[10.38-16.00] | 1.23[1.02-1.47] | 0.033 | **0.072** | 1.19[0.96-1.47] | 0.109 | 0.191 |
| IL2 | 9.00[6.50-11.00] | 11.00[8.00-16.62] | 1.73[0.99-3.02] | 0.055 | 0.101 | 1.45[0.80-2.66] | 0.217 | 0.341 |
| IL6 | 27.00[22.00-32.00] | 23.75[19.38-33.25] | 1.05[0.84-1.31] | 0.685 | 0.754 | 1.00[0.78-1.28] | 0.983 | 0.983 |
| IL13 | 14.00[11.00-18.00] | 19.50[14.00-24.12] | 1.37[1.13-1.67] | 0.002 | **0.010** | 1.30[1.04-1.62] | 0.023 | ***0.076*** |
| IL18 | 27.50[22.00-41.62] | 37.75[27.75-97.38] | 1.29[0.78-2.13] | 0.319 | 0.370 | 1.17[0.67-2.06] | 0.575 | 0.671 |
| TNF | 11.50[10.00-15.00] | 15.00[11.00-24.00] | 1.31[0.99-1.73] | 0.065 | 0.110 | 1.24[0.92-1.69] | 0.158 | 0.258 |
| LTA | 18.50[15.50-22.00] | 21.00[15.75-29.29] | 1.40[1.10-1.77] | 0.007 | **0.022** | 1.36[1.03-1.80] | 0.028 | ***0.076*** |
| **Chemokines** | | | | | | | | |
| CXCL8 | 37.50[29.00-45.00] | 42.25[32.62-75.25] | 1.45[1.19-1.77] | ≤0.001 | **0.003** | 1.41[1.13-1.76] | 0.003 | ***0.020*** |
| CXCL10 | 356.00[176.00-610.00] | 475.00[364.25-660.75] | 1.20[0.89-1.61] | 0.226 | 0.284 | 1.36[0.98-1.32] | 0.063 | 0.126 |
| CCL2 | 267.00[211.50-364.00] | 168.75[98.25-254.50] | 0.73[0.56-0.95] | 0.019 | **0.047** | 0.74[0.54-1.01] | 0.058 | 0.121 |

**Statistics:** Values are expressed as the median [IQR]. P-values estimated using a generalized linear model, adjusted by sex and ethnic origin. Significant values are shown in bold type and were defined as aAMR≥1.2 or ≤ 0.8 and q-value<0.1. **Abbreviations**: PLWH, people living with HIV, HIV, human immunodeficiency virus; HIV/SARS, PLWH previously infected by SARS-CoV-2 (≥ 4 weeks post-infection and diagnosis with PCR+); IQR, interquartile range; aAMR, adjusted arithmetic mean ratio; p, p.value; q, adjusted p.value, p-value corrected for multiple comparisons by Benjamini and Hochberg.

### Table S3. Levels of inflammatory plasma cytokines in PLWH after recovering from SARS-CoV-2 infection.

| | Mean (IQR) | | Univariate analysis | | | Multivariate analysis | | |
|---|---|---|---|---|---|---|---|---|
| | **HIV** | **HIV/SARS** | **AMR** | **p** | **q** | **aAMR** | **p** | **q** |
| **Cytokines** | | | | | | | | |
| IL4 | 11.50[10.00-13.00] | 14.00[11.00-17.00] | 1.34[1.12-1.60] | 0.002 | **0.010** | 1.27[1.03-1.55] | 0.024 | ***0.076*** |
| IL5 | 10.00[8.50-11.00] | 10.50[9.00-13.00] | 1.12[0.98-1.27] | 0.103 | 0.157 | 1.13[0.98-1.32] | 0.102 | 0.188 |
| IL10 | 13.00[10.00-15.00] | 12.75[11.00-16.38] | 1.08[0.92-1.27] | 0.340 | 0.383 | 0.99[0.82-1.20] | 0.936 | 0.957 |
| IL12B | 10.00[8.00-11.00] | 13.25[9.88-17.62] | 1.56[1.26-1.94] | ≤0.001 | **0.002** | 1.57[1.21-2.03] | ≤0.001 | ***0.011*** |
| IL17A | 13.00[11.00-15.00] | 14.50[11.88-19.00] | 1.38[1.16-1.63] | ≤0.001 | **0.003** | 1.30[1.06-1.59] | 0.010 | ***0.038*** |
| **Chemokines** | | | | | | | | |
| CCL3 | 42.00[32.00-56.00] | 56.25[42.12-102.75] | 2.62[1.64-4.18] | ≤0.001 | **0.002** | 2.60[1.47-4.61] | ≤0.001 | ***0.011*** |
| CCL4 | 82.00[66.00-115.00] | 98.00[73.75-139.50] | 1.19[0.99-1.44] | 0.073 | 0.120 | 1.25[1.01-1.56] | 0.042 | ***0.098*** |
| **Interferons** | | | | | | | | |
| IFNA1 | 22.50[20.00-26.00] | 23.25[19.38-31.00] | 1.17[1.02-1.34] | 0.026 | 0.061 | 1.19[1.02-1.40] | 0.027 | ***0.076*** |

**Statistics:** Values are expressed as the median [IQR]. P-values estimated using a generalized linear model, adjusted by sex and ethnic origin. Significant values are shown in bold type and were defined as aAMR≥1.2 or ≤ 0.8 and q-value<0.1. **Abbreviations**: PLWH, people living with HIV, HIV, human immunodeficiency virus; HIV/SARS, PLWH previously infected by SARS-CoV-2 (≥ 4 weeks post-infection and diagnosis with PCR+); IQR, interquartile range; aAMR, adjusted arithmetic mean ratio; p, p.value; q, adjusted p.value, p-value corrected for multiple comparisons by Benjamini and Hochberg.

**Table S4. Immune checkpoint-related plasma biomarkers in PLWH after recovering from SARS-CoV-2 infection.**

| | | Mean (IQR) | | Univariate analysis | | | Multivariate analysis | | |
|---|---|---|---|---|---|---|---|---|---|
| | | HIV | HIV/SARS | AMR | p | q | aAMR | p | q |
| **Immunoglobuline superfamily** | | | | | | | | | |
| B7 | *CD80* | 26.00[21.12-32.00] | 36.00[29.75-71.00] | 1.64[1.19-2.25] | 0.003 | **0.012** | 1.80[1.27-2.56] | 0.001 | **0.011** |
| | *ICOSL* | 270.50[222.00-305.00] | 279.75[227.75-345.88] | 1.08[0.97-1.21] | 0.175 | 0.233 | 1.07[0.95-1.22] | 0.282 | 0.401 |
| | *CD274* | 18.50[16.00-22.00] | 19.25[16.62-22.50] | 1.24[0.89-1.71] | 0.199 | 0.258 | 1.21[0.87-1.67] | 0.273 | 0.400 |
| | *PDCD1LG2* | 91.50[77.00-106.88] | 113.00[92.50-142.00] | 1.27[1.09-1.48] | 0.002 | **0.010** | 1.25[1.06-1.48] | 0.010 | **0.038** |
| | *NCR3LG1* | 16.50[14.00-20.50] | 17.00[15.00-21.25] | 0.98[0.80-1.20] | 0.829 | 0.862 | 0.94[0.75-1.18] | 0.579 | 0.671 |
| | *CD276* | 3020.0[2393.50-3668.25] | 3888.75[3209.62-4536.62] | 1.27[1.13-1.42] | 0.000 | **0.002** | 1.28[1.13-1.45] | 0.000 | **0.011** |
| | *VSIR* | 16.00[13.25-21.00] | 18.00[15.00-24.62] | 1.39[0.95-2.03] | 0.095 | 0.149 | 1.20[0.85-1.72] | 0.304 | 0.406 |
| CD28 | *CD28* | 20.25[18.00-24.00] | 26.00[20.00-44.00] | 1.24[0.81-1.89] | 0.318 | 0.370 | 1.13[0.71-1.82] | 0.607 | 0.675 |
| | *PDCD1* | 29.50[24.00-40.38] | 41.00[33.00-63.75] | 1.61[1.23-2.10] | 0.001 | **0.005** | 1.65[1.24-2.20] | 0.001 | **0.011** |
| | *BTLA* | 62.25[44.38-82.25] | 69.5[057.50-94.50] | 1.29[0.92-1.82] | 0.140 | 0.198 | 1.32[0.92-1.90] | 0.127 | 0.215 |
| | *CTLA4* | 28.50[24.00-44.00] | 39.00[28.75-56.50] | 1.56[0.99-2.46] | 0.057 | 0.101 | 1.18[0.79-1.80] | 0.421 | 0.529 |
| CD2 | *CD48* | 14.00[11.50-16.00] | 17.00[13.38-22.00] | 1.34[1.06-1.68] | 0.015 | **0.041** | 1.16[0.91-1.48] | 0.226 | 0.342 |
| Other | *CD47* | 24.00[19.50-29.25] | 26.50[22.00-34.00] | 1.16[1.01-1.34] | 0.036 | 0.075 | 1.25[1.08-1.46] | 0.004 | **0.023** |
| | *LAG3* | 19.50[16.25-24.38] | 25.50[21.00-32.50] | 1.06[0.77-1.45] | 0.727 | 0.780 | 1.09[0.75-1.57] | 0.662 | 0.710 |
| **TIM family** | | | | | | | | | |
| *HAVCR2* | | 145.00[114.12-169.75] | 154.75[131.75-192.75] | 1.18[1.05-1.33] | 0.007 | 0.023 | 1.25[1.10-1.42] | 0.001 | **0.011** |
| *TIMD4* | | 125.00[79.75-191.00] | 149.00[98.25-251.50] | 1.22[0.95-1.56] | 0.125 | 0.184 | 1.37[1.03-1.82] | 0.030 | **0.078** |
| **TNF/RSF family** | | | | | | | | | |
| *TNFRSF9* | | 34.25[28.00-45.38] | 45.00[33.50-66.75] | 1.34[1.00-1.80] | 0.052 | 0.101 | 1.39[1.03-1.87] | 0.034 | **0.082** |
| *CD27* | | 2958.50[2348.25-3869.50] | 2630.50[1478.50-3648.75] | 0.84[0.70-1.00] | 0.056 | 0.101 | 0.90[0.73-1.10] | 0.301 | 0.406 |
| *TNFRSF18* | | 23.00[21.00-28.75] | 28.50[21.75-43.00] | 1.53[1.08-2.15] | 0.018 | 0.047 | 1.66[1.16-2.37] | 0.007 | **0.033** |
| *TNFRSF14* | | 25.00[22.00-27.8] | 31.00[25.12-36.75] | 1.46[1.20-1.77] | 0.000 | **0.003** | 1.35[1.09-1.69] | 0.007 | **0.033** |
| *TNFRSF4* | | 35.00[29.00-39.75] | 39.50[34.00-49.50] | 1.37[1.08-1.75] | 0.012 | **0.036** | 1.30[1.01-1.69] | 0.046 | 0.101 |
| **S100 family** | | | | | | | | | |
| *S100A8A9* | | 73.00[55.25-106.50] | 75.75[66.00-95.75] | 0.99[0.86-1.14] | 0.901 | 0.901 | 1.01[0.86-1.19] | 0.867 | 0.908 |
| **Enzyme** | | | | | | | | | |
| *IDO* | | 45.50[28.25-90.38] | 98.00[55.00-246.50] | 3.10[1.40-6.84] | 0.006 | **0.021** | 2.04[0.92-4.60] | 0.066 | 0.126 |

**Statistics:** Values are expressed as the median [IQR]. P-values estimated using a generalized linear model, adjusted by sex and ethnic origin. Significant values are shown in bold type and were defined as aAMR≥1.2 or ≤ 0.8 and q-value<0.1. **Abbreviations**: PLWH, people living with HIV, HIV, human immunodeficiency virus; HIV/SARS, PLWH previously infected by SARS-CoV-2 (≥ 4 weeks post-infection and diagnosis with PCR+); IQR, interquartile range; aAMR, adjusted arithmetic mean ratio; p, p.value; q, adjusted p.value, p-value corrected for multiple comparisons by Benjamini and Hochberg.

**Table S5. Relationship between significant plasma biomarkers in HIV/SARS patients.**

| | | Immune checkpoint | | | | | | | | | | | | | | | | | | | |
|---|---|---|---|---|---|---|---|---|---|---|---|---|---|---|---|---|---|---|---|---|---|
| | | **CD80** | | **PDCD1LG2** | | **CD276** | | **PDCD1** | | **CD47** | | **HAVCR2** | | **TIMD4** | | **TNFRSF9** | | **TNFRSF18** | | **TNFRSF14** | |
| | | Rho | q | Rho | q | Rho | q | Rho | q | Rho | q | Rho | q | Rho | q | Rho | q | Rho | q | Rho | q |
| **SASP** | **IL13** | 0.31 | ***0.073**** | -0.20 | 0.600 | -0.34 | 0.151 | 0.24 | 0.256 | -0.12 | 0.852 | -0.12 | 0.678 | 0.07 | 0.809 | 0.26 | 0.173 | 0.37 | ***0.024**** | 0.33 | ***0.071**** |
| | **LTA** | 0.34 | ***0.073**** | -0.11 | 0.600 | -0.27 | 0.188 | 0.31 | 0.152 | -0.11 | 0.852 | -0.11 | 0.678 | 0.10 | 0.809 | 0.25 | 0.173 | 0.44 | ***0.019**** | 0.46 | ***0.012**** |
| | **CXCL8** | 0.23 | 0.174 | -0.25 | 0.600 | -0.07 | 0.731 | 0.07 | 0.662 | 0.02 | 0.905 | -0.15 | 0.678 | 0.09 | 0.809 | 0.10 | 0.494 | 0.19 | 0.229 | 0.09 | 0.697 |
| **Pro and anti-inflammatory** | **IL4** | 0.19 | 0.247 | -0.10 | 0.600 | -0.08 | 0.731 | 0.15 | 0.423 | -0.08 | 0.852 | -0.16 | 0.678 | -0.05 | 0.809 | 0.13 | 0.442 | 0.33 | ***0.040**** | 0.07 | 0.697 |
| | **IL12B** | 0.31 | ***0.073**** | -0.10 | 0.600 | -0.31 | 0.151 | 0.23 | 0.256 | 0.13 | 0.852 | -0.05 | 0.810 | 0.04 | 0.809 | 0.33 | 0.173 | 0.29 | 0.070 | 0.30 | 0.098 |
| | **IL17A** | 0.10 | 0.562 | -0.17 | 0.600 | -0.16 | 0.613 | 0.18 | 0.346 | -0.05 | 0.852 | -0.06 | 0.810 | 0.08 | 0.809 | 0.25 | 0.173 | 0.23 | 0.164 | 0.34 | ***0.071**** |
| | **CCL3** | 0.27 | 0.128 | -0.09 | 0.600 | -0.08 | 0.731 | 0.22 | 0.256 | 0.20 | 0.852 | -0.02 | 0.915 | 0.15 | 0.809 | 0.23 | 0.173 | 0.37 | ***0.024**** | 0.19 | 0.302 |
| | **CCL4** | 0.03 | 0.843 | 0.03 | 0.862 | 0.01 | 0.949 | 0.12 | 0.482 | -0.07 | 0.852 | 0.15 | 0.678 | 0.16 | 0.809 | 0.17 | 0.316 | 0.13 | 0.406 | 0.06 | 0.697 |
| | **INFA1** | 0.32 | ***0.073**** | -0.10 | 0.600 | -0.08 | 0.731 | 0.33 | 0.152 | -0.06 | 0.852 | -0.21 | 0.678 | -0.08 | 0.809 | 0.24 | 0.173 | 0.39 | ***0.024**** | 0.22 | 0.239 |

**Statistics:** Rho, Spearman's rank correlation coefficient; q, p.value estimated using a Spearman correlation and adjusted by false discovery rate (FDR) using Benjamin-Hochberg correction setting a cut-off point of 0.1. ***** Significant values significant values are shown in bold and italics and were defined as Rho≥0.3 and q<0.1. **Abbreviations**: HIV, human immunodeficiency virus; SASP, senescence-associated secretory phenotype.

**Table S6. Levels of plasma biomarkers in PLWH according to the severity of SARS-CoV-2 infection.**

| | Median (IQR) | | Univariate analysis | | |
|---|---|---|---|---|---|
| | **Mild** | **Severe** | **AMR** | **p** | **q** |
| CSF3 | 16.00[13.50-19.00] | 16.50[15.50-18.25] | 0.97[0.77-1.24] | 0.808 | 0.947 |
| CSF1 | 10.75[9.00-14.00] | 8.50[7.75-9.50] | 0.70[0.47-1.08] | ≤0.001 | 0.741 |
| IFNA1 | 24.00[21.00-31.75] | 19.50[16.75-25.38] | 0.81[0.60-1.12] | 0.188 | 0.741 |
| IFNG | 31.00[20.00-62.00] | 27.50[18.88-93.50] | 1.32[0.68-2.90] | 0.451 | 0.947 |
| IL1B | 13.50[10.88-16.00] | 11.00[9.75-13.38] | 0.91[0.69-1.22] | 0.524 | 0.947 |
| IL10 | 12.75[11.00-18.00] | 12.50[11.00-14.50] | 0.90[0.68-1.21] | 0.477 | 0.947 |
| IL12B | 13.75[10.00-17.62] | 12.50[9.25-16.12] | 0.97[0.59-1.70] | 0.917 | 0.947 |
| IL13 | 19.50[14.75-24.00] | 18.75[10.25-27.12] | 0.90[0.59-1.45] | 0.658 | 0.947 |
| IL17A | 14.00[11.38-18.38] | 15.50[12.38-19.00] | 0.94[0.65-1.43] | 0.774 | 0.947 |
| IL18 | 37.75[27.75-90.75] | 38.00[28.00-129.75] | 0.92[0.44-2.24] | 0.841 | 0.947 |
| IL2 | 11.00[9.75-17.25] | 8.50[8.00-11.00] | 0.47[0.18-1.52] | 0.156 | 0.741 |
| IL4 | 14.00[12.00-17.00] | 9.75[8.00-16.25] | 0.88[0.60-1.34] | 0.539 | 0.947 |
| IL5 | 10.50[9.00-13.62] | 10.50[8.00-11.62] | 0.85[0.65-1.13] | 0.250 | 0.741 |
| IL6 | 26.00[19.00-34.62] | 22.00[20.25-25.25] | 0.76[0.49-1.24] | 0.256 | 0.741 |
| CXCL8 | 42.75[32.2-81.25] | 40.00[33.62-60.50] | 0.91[0.59-1.49] | 0.705 | 0.947 |
| CXCL10 | 466.75[356.50-707.75] | 556.50[396.12-649.75] | 0.95[0.67-1.39] | 0.780 | 0.947 |
| CCL2 | 161.50[88.50-253.00] | 221.75[164.50-283.25] | 1.37[0.78-2.58] | 0.304 | 0.776 |
| CCL3 | 56.75[42.88-105.88] | 51.75[35.50-75.25] | 1.51[0.19-1.77] | 0.230 | 0.741 |
| CCL4 | 98.00[70.25-142.50] | 98.00[81.75-117.00] | 0.97[0.67-1.44] | 0.869 | 0.947 |
| TNF | 15.50[12.00-24.00] | 13.00[9.75-19.50] | 1.01[0.70-1.53] | 0.944 | 0.947 |
| LTA | 22.00[17.38-31.12] | 15.00[14.75-19.00] | 0.62[0.37-1.10] | ≤0.001 | 0.741 |
| BTLA | 70.00[57.50-98.50] | 65.00[54.75-73.00] | 1.17[0.60-2.54] | 0.671 | 0.947 |
| TNFRSF9 | 46.00[33.00-72.75] | 42.50[36.00-51.75] | 1.11[0.69-1.87] | 0.686 | 0.947 |
| CTLA4 | 39.00[28.75-56.50] | 42.50[28.75-71.38] | 1.87[0.83-5.05] | 0.174 | 0.741 |
| CD27 | 2630.50[1546.75-3648.75] | 2957.25[999.75-3580.50] | 0.96[0.65-1.46] | 0.838 | 0.947 |
| CD28 | 25.50[20.00-43.50] | 28.75[23.00-67.25] | 1.60[1.03-2.60] | ≤0.001 | 0.598 |
| CD80 | 36.00[30.25-71.00] | 41.25[29.25-57.00] | 0.89[0.51-1.67] | 0.697 | 0.947 |
| TNFRSF18 | 32.50[22.50-43.50] | 23.75[19.88-28.88] | 0.65[0.32-1.53] | 0.285 | 0.772 |
| TNFRSF14 | 31.00[25.00-37.75] | 33.00[29.50-34.12] | 0.84[0.54-1.38] | 0.476 | 0.947 |
| IDO1 | 103.00[56.00-229.75] | 83.00[42.12-724.62] | 1.68[0.42-11.98] | 0.525 | 0.947 |
| LAG3 | 25.00[21.00-31.00] | 33.25[24.88-44.88] | 1.49[1.16-1.95] | ≤0.001 | ***≤0.001**** |
| PDCD1 | 46.00[33.75-67.00] | 37.50[30.38-41.00] | 0.73[0.44-1.29] | 0.258 | 0.741 |
| CD274 | 19.75[16.62-22.50] | 18.25[17.25-22.62] | 1.49[0.79-3.11] | 0.258 | 0.741 |
| PDCD1LG2 | 113.00[94.75-138.50] | 109.25[88.62-142.50] | 1.02[0.78-1.34] | 0.907 | 0.947 |
| HAVCR2 | 154.75[128.62-193.75] | 155.00[140.88-180.00] | 0.96[0.76-1.22] | 0.712 | 0.947 |
| NCR3LG1 | 17.00[15.00-22.00] | 18.00[15.75-20.62] | 0.95[0.76-1.20] | 0.669 | 0.947 |
| TNFRSF4 | 39.50[33.75-48.25] | 38.75[36.25-59.12] | 1.02[0.57-2.00] | 0.947 | 0.947 |
| CD276 | 3805.75[3209.62-4569.25] | 4183.25[3272.38-4477.75] | 0.99[0.83-1.20] | 0.934 | 0.947 |
| CD47 | 27.00[22.00-34.00] | 25.50[22.12-28.50] | 0.91[0.68-1.25] | 0.564 | 0.947 |
| CD48 | 17.50[13.38-22.00] | 14.75[13.75-17.50] | 0.74[0.47-1.23] | 0.229 | 0.741 |
| ICOSL | 279.75[223.88-352.50] | 293.75[253.75-343.00] | 1.03[0.82-1.32] | 0.793 | 0.947 |
| S100A8/A9 | 79.50[68.50-99.00] | 57.75[52.88-66.75] | 0.74[0.61-0.90] | ≤0.001 | ***≤0.001**** |
| TIMD4 | 147.50[87.25-233.25] | 204.00[135.88-270.75] | 1.29[0.80-2.18] | 0.324 | 0.784 |
| VSIR | 18.00[14.88-25.25] | 17.50[15.75-19.12] | 0.74[0.32-2.13] | 0.538 | 0.947 |

**Statistics**: Values are expressed as the median [IQR]. P-values estimated using a generalized linear model, adjusted by sex and etnia. *****Significant values significant values are shown in bold and italics and were defined as aAMR≥1.2 or ≤ 0.8 and q-value<0.1. **Abbreviations:** PLWH, people living with HIV; IQR, interquartile range; AMR, arithmetic mean ratio; p, p.value; q, p-value corrected for multiple comparisons by Benjamini and Hochberg.
